# Inactivation of thermogenic UCP1 as a historical contingency in multiple placental mammal clades

**DOI:** 10.1101/086819

**Authors:** Michael J. Gaudry, Martin Jastroch, Jason R. Treberg, Michael Hofreiter, Johanna L.A. Paijmans, James Starrett, Nathan Wales, Anthony V. Signore, Mark S. Springer, Kevin L. Campbell

## Abstract

Mitochondrial uncoupling protein 1 (UCP1) is essential for non-shivering thermogenesis in brown adipose tissue and is widely accepted to have played a key thermoregulatory role in small-bodied and neonatal placental mammals that enabled the exploitation of cold environments. Here we map *ucp1* sequences from 133 mammals onto a species tree constructed from a ∼51-kb sequence alignment and show that inactivating mutations have occurred in at least eight of the 18 traditional placental orders, thereby challenging the physiological importance of UCP1 across Placentalia. Selection and timetree analyses further reveal that *ucp1* inactivations temporally correspond with strong secondary reductions in metabolic intensity in xenarthrans and pangolins, or in six other lineages coincided with a ∼30 million year episode of global cooling in the Paleogene that promoted sharp increases in body mass and cladogenesis evident in the fossil record. Our findings also demonstrate that members of various lineages (*e.g*., cetaceans, horses, woolly mammoths, Steller’s sea cows) evolved extreme cold hardiness in the absence of UCP1-mediated thermogenesis. Finally, we identify *ucp1* inactivation as a historical contingency that is linked to the current low species diversity of clades lacking functional UCP1, thus providing the first evidence for species selection related to the presence or absence of a single gene product.

## INTRODUCTION

Adaptive non-shivering thermogenesis (NST) of placental mammals is predominantly mediated by uncoupling protein 1 (UCP1), which resides at high levels in the mitochondrial inner membrane of brown adipose tissue (BAT) (1). Upon activation, UCP1 facilitates proton leak into the mitochondrial matrix (2) without concomitant ATP production, thereby intensifying substrate oxidation and hence cellular heat production. BAT deposits are located adjacent to major blood vessels in the neck, thorax, and abdomen to promote the rapid distribution of NST heat (3). In addition to its strategic location, the observation that chronic cold stress stimulates the proliferation and activity of BAT (4), highlights the canonical importance of NST for small-bodied and torpor expressing placental mammals, together with neonates of larger cold-tolerant species (5). UCP1 has also been implicated in the regulation of food intake, energy balance, the fever response, the prevention of cold-induced oxidative stress, and possibly even longevity (6-9). The recent discovery of UCP1 induction in a class of white adipocytes (termed brite or beige fat) in response to exercise and cold exposure expands the repertoire of this protein and moreover offers therapeutic interventions for obesity, hyperlipidemia, cachexia, and other metabolic disorders (10, 11).

The presence of non-thermogenic UCP1 in marsupials, monotremes, as well as both ray-finned and lobe-finned fishes (12) indicates that *ucp1* was likely recruited for NST sometime during the Mesozoic ancestry of placental mammals (13, 14). This ancestor is reconstructed as a small (6 to 245 g) nocturnal insectivore that gave birth to hairless, altricial young and likely exhibited torpor (*i.e*., heterothermy) (15, 16). Facultative BAT thermogenesis may have evolved to defend against the natural cold stress of birth (17), to decrease the energetic costs of rewarming from torpor (18), or to enable endothermy during the reproductive season (19), and is postulated to underlie the subsequent radiation of crown eutherians into cold environments (3). Evolution of this thermogenic tissue was presumably also driven by small body size as maximal NST capacity is inversely proportional to mass, with little to no thermal contribution predicted in mammals above 10 kg (20, 21). Although limited data are available for larger bodied species, *ucp1* is only transiently expressed in BAT deposits of neonatal cattle (∼40 kg; 22) and pre-weaned harp seals (<30 kg; *23*), with minor depots either absent or present in adult humans (11). Conversely, the persistence of UCP1 in BAT depots of subadult harbor seals (34–38 kg; 5) and the discovery of a large cervical mass of fat that is histologically similar to BAT in a preserved 1-month old (∼100 kg) woolly mammoth (24), but not extant elephants, implies this tissue may underpin the evolution of extreme cold hardiness in at least some larger-bodied species. Consistent with this hypothesis, *ucp1* inactivation is only documented within the pig lineage whose newborn lack discernible BAT and are notoriously cold intolerant (17, 25).

Support for a *de novo* gain of thermogenic function in early eutherians is bolstered by an elevated rate of nucleotide substitutions on the stem placental branch (13, 14), though these results were based on limited sample sizes (10 and 16 species, respectively) and heavily biased towards small-bodied forms from only two (Euarchontoglires and Laurasiatheria) of the four major clades of placental mammals. Notable in this regard, distinct BAT deposits have only been positively identified in neonates of 11 of the 18 traditional eutherian orders (17, 26). Given that body size and energetic considerations are expected to alter the selective advantage of BAT-mediated NST (20, 21, *26*), we posited that evolution of large body size or reduced thermogenic capacity would be accompanied by relaxed selection and/or inactivation of the *ucp1* locus that may, at least in part, underlie this latter observation. To test this hypothesis, we employed genome mining, polymerase chain reaction, and hybridization capture techniques to obtain *ucp1*, *ucp2*, and *ucp3* coding sequences from 141 vertebrate species (see Materials and Methods; table S1). Translated coding sequences were examined for disrupting mutations (*e.g*., frameshift indels, premature termination codons, mutated start/stop codons, and splice site mutations), and ortholog assignments verified on the basis of synteny analyses and/or a maximum likelihood *ucp1*-*ucp2*-*ucp3* tree (fig. S1) prior to conducting selection analyses using the CODEML program in the PAML 4.8 software package (27) based on a custom built 41-loci (51-kb) RAxML species tree (see Materials and Methods; data file S1).

## RESULTS

In line with previous studies (13, 14), our phylogenetically informed results demonstrate an increased rate of molecular evolution on the stem placental branch for *ucp1* (ω=0.65) that was presumably associated with the acquisition of classical NST, though no sites exhibited statistically significant signatures of positive selection (table S2). These analyses further verify that eutherian branches with intact *ucp1* sequences evolved under strong purifying selection (ω=0.16), consistent with maintenance of an adaptive function. Importantly, in support of our hypothesis, these analyses reveal independent inactivating mutations of *ucp1* (including exon and whole gene deletions) in at least eight placental clades: xenarthrans (Cingulata and Pilosa), pangolins (Pholidota), dolphins and whales (Cetacea), sea cows (Sirenia), elephants (Proboscidea), hyraxes (Hyracoidea), pigs (Suidae), and horses, donkeys, and zebras (Equidae) (Figs. 1, 2 and fig. S2). Notably, these findings are corroborated by a signature of neutral evolution (ω=0.94 in pseudogenic branches versus 0.16 in species containing an intact *ucp1* locus; table S2 and data file S2) and a lack of discernable BAT in neonates of each of these groups (17). To better facilitate comparisons between the genomic record and fossil record, and identify potential climatic and ecological factors underlying *ucp1* loss in these lineages, we calculated gene inactivation date ranges based on a fossil-informed Bayesian molecular timetree that employed an autocorrelated rates model (see Materials and Methods; data file S1), and d*N*/d*S* ratios in functional, pseudogenic, and mixed branches (where *ucp1* inactivation was deemed to have occurred; *28*).

**Fig. 1.**
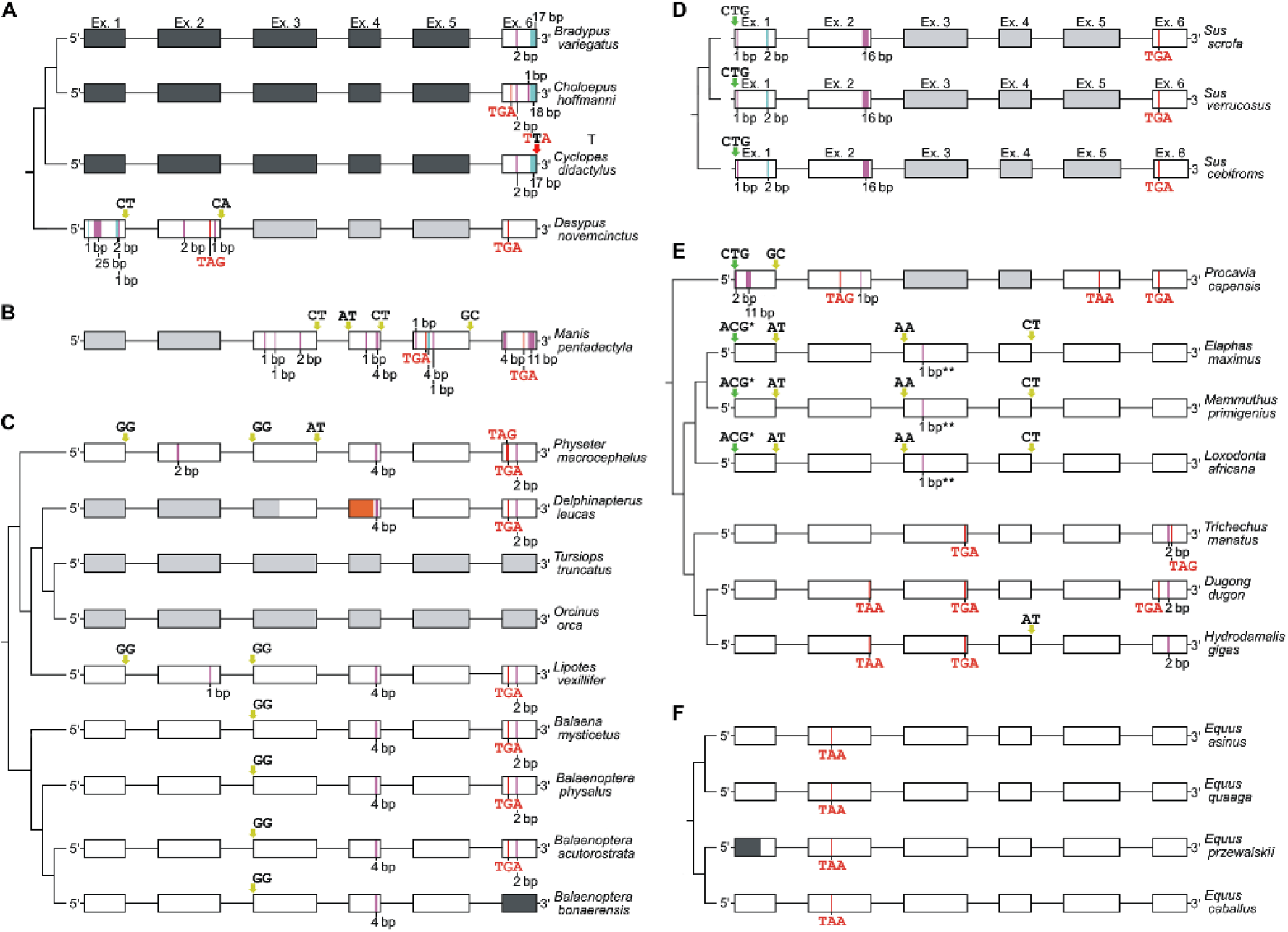
Schematic illustrating disrupting mutations in *ucp1* across the placental mammal phylogeny. (**A**) Xenarthra, (**B**) Pholidota, (**C**) Cetacea; note insert from ∼800 bp upstream of *ucp1* exon 4 of beluga (orange), (**D**) Suidae, (**E**) Paenungulata; *putative start codon located 36 bp downstream, **creates 15 downstream stop codons (not shown), (**F**) Equidae. The six coding exons of *ucp1* are represented by open rectangles; missing data (black), deleted exons (grey), mutated start codons (green), nonsense mutations (red), deletions (magenta), insertions (cyan), splice site mutations (yellow), and a mutated stop codon (‘TTA’ in *Cyclopes*) are indicated.

**Fig. 2.**
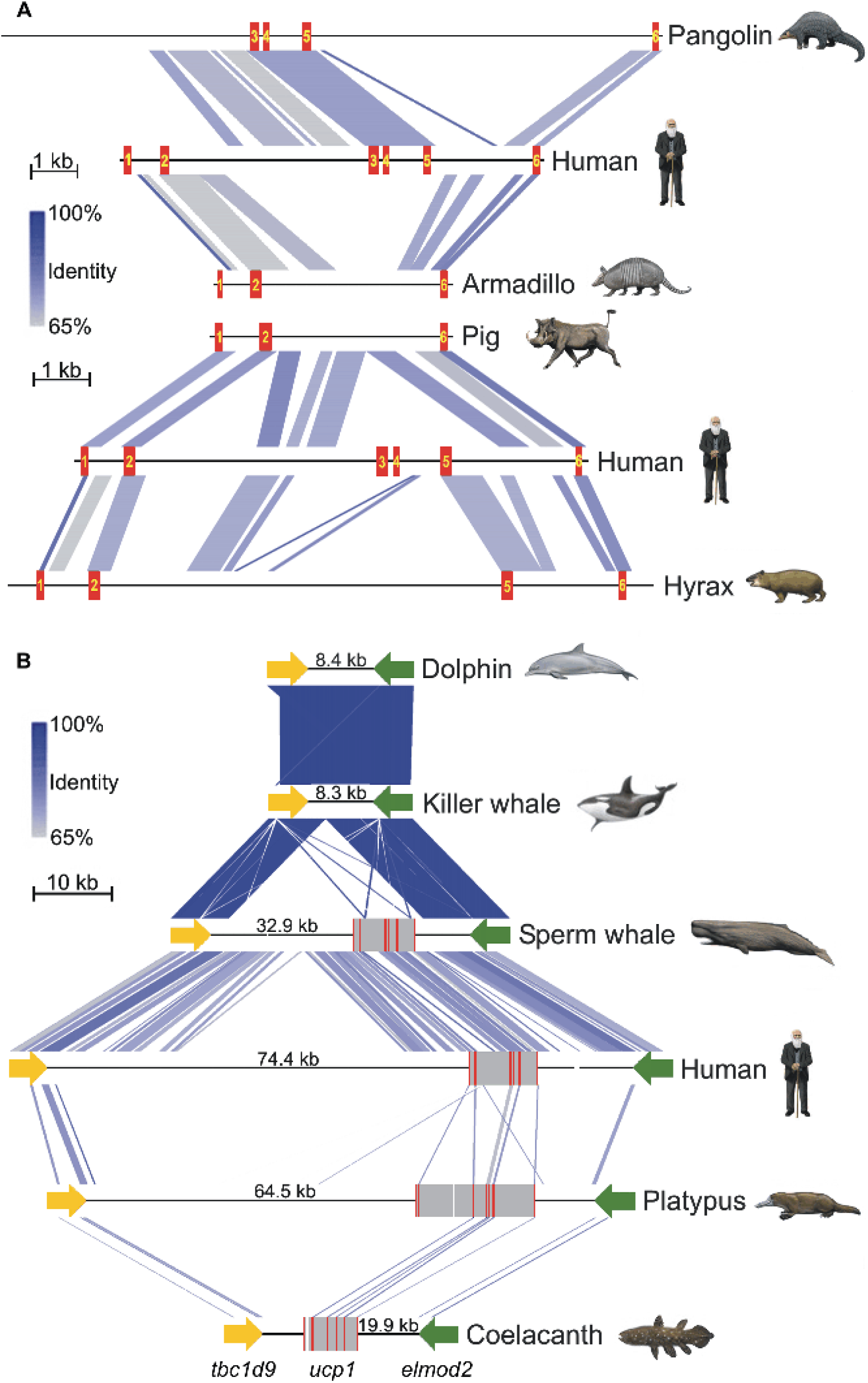
Large scale deletions of the uncoupling protein 1 (*ucp1*) locus in select placental species. (**A**) Linear comparisons of *ucp1* illustrating exon deletions in pangolin, armadillo, pig, and hyrax relative to the intact human locus. The coding exons of *ucp1* are numbered and highlighted in red. (**B**) Patterns of conserved synteny in the genomic region flanking the *ucp1* locus of select vertebrates illustrating the complete deletion of *ucp1* in a common ancestor of the dolphin and killer whale. Coding exons are highlighted in red, intervening sequences are highlighted in grey, while sequencing gaps are denoted by white spaces. Distances in kilobases (kb) between the termination codons of the upstream (*tbc1d9*) and downstream (*elmod2*) loci are given for each species. Note that the platypus *ucp1* locus contains an additional upstream exon relative to other vertebrate species. Artwork by Carl Buell.

The earliest disruptions of *ucp1* function (likely Cretaceous; Fig. 3 and table S3) occur in the ancestors of xenarthrans and pangolins, two groups that exhibit remarkable convergence in many ecological, morphological, and physiological traits and which are generally characterized by ‘inferior’ temperature-regulating mechanisms, and low, labile body temperatures (32-35). The ancient inactivations of *ucp1* in these lineages may in part underlie this convergence and are presumably linked to strong secondary reductions in metabolic intensity associated with the ancestral burrowing habits and energy poor diets of these taxa (32, 33). While aardvarks share some ecological and dietary commonalities with pangolins and insectivorous xenarthrans, they only exhibit minor reductions in mass-specific basal metabolic rate (∼20% lower than expected for its body mass) compared to pangolins, armadillos, and anteaters of similar size (∼50-65% lower than predicted) (36), and are hence likely able to adequately provision offspring with thermogenic BAT (*i.e.*, with elevated energetic costs). Interestingly, while BAT-mediated NST has been shown to enhance cold tolerance (37) and accelerate rewarming from torpor at a reduced net energy cost (18), at least some xenarthrans exhibit torpor (35) while others exploited thermally taxing environments. The latter was presumably enabled by elaborate countercurrent heat exchangers (arteriovenus rete) in the limbs that promote heat conservation (38) which, coupled with a relatively thick pelt and the recurrent development of gigantism (39), likely facilitated the colonization of subarctic environments (68°N) by the Pleistocene sloth *Megalonyx jeffersonii* (40) and the aquatic niche by Miocene/Pliocene marine sloths of the genus *Thalassocnus* (41) in the absence of functional UCP1.

**Fig. 3.**
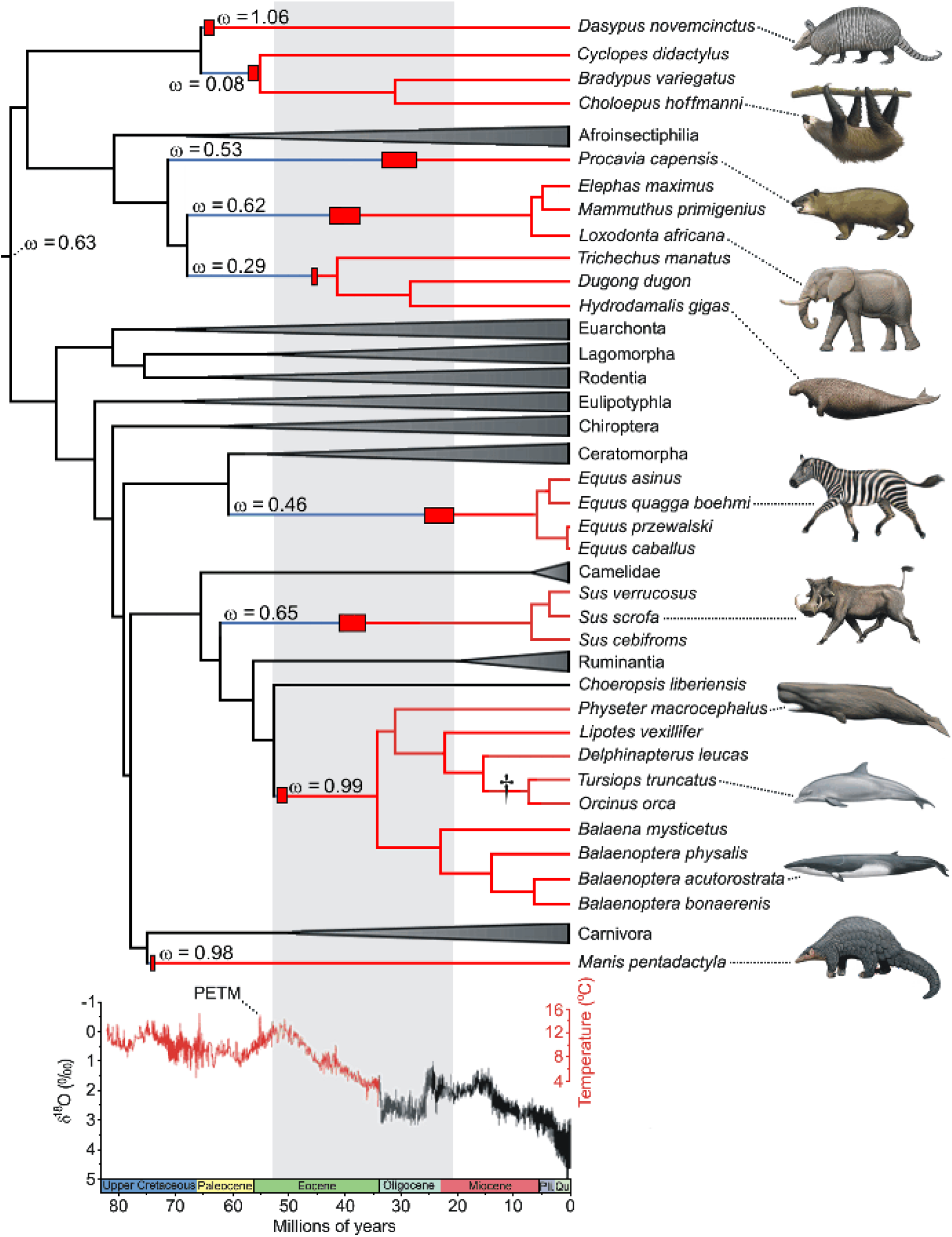
UCP1 inactivation across a time-calibrated placental mammal phylogeny. Functional *ucp1* branches are denoted in black, pseudogenic branches in red, and mixed branches (where *ucp1* inactivation occurred) in blue. Clades possessing intact *ucp1* loci are collapsed. Red boxes indicate *ucp1* inactivation date ranges as determined by d*N*/d*S* analyses (see table S3), while the dagger represents the complete deletion of this locus in the delphinid lineage (see Fig. 2B). Pre-Oligocene temperatures are based upon a benthic foraminifera δ^18^O isotope dataset assuming an ice-free ocean (29-31). Note that the majority of inactivations correspond to a period of intense global cooling (grey shading) following the Paleocene-Eocene Thermal Maximum (PETM). Within this window, *ucp1* was inactivated earlier in the two aquatic species, consistent with the much higher thermal conductivity of water relative to air (∼24.1× higher at 25°C). As only remnants of *ucp1* were identified from representative Pilosa (*Bradypus*, *Choleopus*, and *Cyclopes*), the presented inactivation range should be interpreted with caution, and it is possible that *ucp1* pseudogenization preceded its divergence from Cingulata. Artwork by Carl Buell.

For the remaining six lineages, our estimates indicate *ucp1* was predominantly silenced from the early Eocene to the late Oligocene, a period characterized by marked global cooling (Fig. 3) that spurred the diversification of grass and deciduous woodland communities (42). As these factors have been proposed to underlie body mass increases of numerous mammalian groups during the Cenozoic (43), we assembled a dataset to explore potential links between mammalian body size evolution (data file S3) and *ucp1* inactivation (table S3). In each case, our *ucp1* inactivation date ranges overlap or immediately precede magnitude scale increases in body size (Fig. 4A). For example, the earliest proboscideans (59-60 million year old [Ma] *Eritherium*; *44*) and hyraxes (∼55 Ma *Seggeurius*; 45) were small (∼3-5 kg), but gave rise to much larger species (*e.g*., 3600 kg *Barytherium* and 800–1400 kg *Titanohyrax ulitimus*) coincident with our estimates of *ucp1* inactivation. Although large-bodied (∼625–800 kg) hyracoids such as ‘*Titanohyrax*’ *mongereaui* are also known from earlier deposits, the presence of only a single, ‘primitive’ *Seggeurius*-sized species (*Dimaitherium patnaiki*) from the early Priabonian (∼37 Ma), suggests that gigantism evolved independently in mid- and late-Eocene hyracoids (46). While extant hyraxes are generally rabbit-sized (∼1–5 kg), and thought to have arisen from small, derived saghatheriines such as *Thyrohyrax* (45), their long gestation (8 months) (45) coupled with multiple disruptions in *ucp1* (Figs. 1 and 2A) is instead suggestive of evolutionary descent from a large-bodied ancestor. Our estimate of *ucp1* inactivation in the family Equidae (25.1–20.7 Ma) is also coincident with an ∼8-10 fold increase in body size in this lineage (Fig. 4A). Potential thermogenic limitations arising from UCP1 loss (as suggested for suids; *25*) were clearly no impediment to the ability of equines to exploit cold environments given “that Alaskan horses prospered during the LGM [last glacial maximum; ∼18,000 years ago], and appear to have been particularly well adapted to the more intense versions of the cold/arid Mammoth Steppe” (47).

**Fig. 4.**
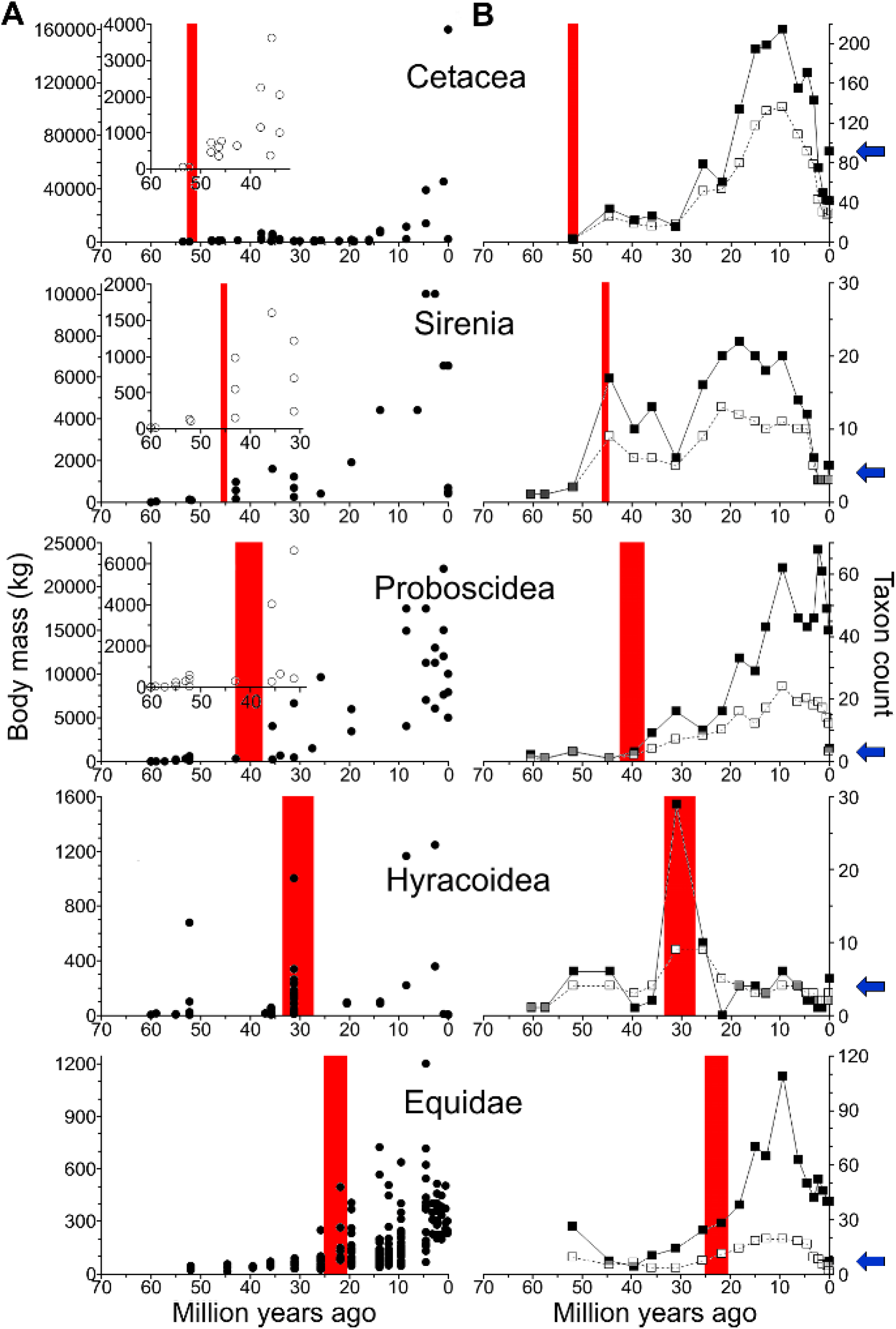
Body mass and taxon diversity relative to estimates of *ucp1* inactivation in five lineages of eutherian mammals. (**A**) Calculated *ucp1* inactivation range (red shading) in relation to body size estimates (open and closed circles). The y-axis is reduced in the insets to better visualize changes in body mass relative to *ucp1* inactivation. (**B**) Calculated *ucp1* inactivation range (red shading) in relation to records of the number of species (closed squares) and genera (open squares) per geological stage. Blue arrows denote current species diversity for each taxon.

Loss of *ucp1* in pigs has been previously dated to ∼20 Ma (25). However, the absence of BAT in newborn peccaries (17) suggests that its inactivation may have predated the divergence of Suoidea in the late Eocene to early Oligocene (48, 49). Owing to a fragmentary fossil record, potential selection pressures favoring *ucp1* inactivation in early suids remain unclear, though it is notable that—like whales and adult pinnipeds—cold-stressed domestic swine shunt blood away from the entire body surface thereby forming a steep thermal gradient to minimize heat loss (50). Nonetheless, loss of BAT has not prevented exploitation of cold north-temperature niches by some members of this clade (25).

We estimate *ucp1* inactivation in Cetacea occurred soon after their divergence from hippopotamids (∼52.75 Ma), and document the complete deletion of this locus in delphinids (Fig. 2B) by the late Miocene. These findings directly contradict recent claims of UCP1 expression in blubber of four delphinoid species (51) (which we attribute to unspecific antibody hybridization to UCP1 paralogs in this tissue), and intact UCP1 proteins in the genomes of six cetacean species (including bottlenose dolphin and killer whale; 52). Notably, the non-delphinid cetacean UCP1 primary sequences presented in the latter study show premature stop codons stemming from both nonsense and frameshift mutations (as also seen in Fig. 1), while the presented killer whale and bottlenose dolphin ‘UCP1’ sequences correspond to UCP2 and UCP3, respectively. Our results further indicate that the pseudogenization of *ucp1* also immediately predates strong increases in body size during the early amphibious stages of archaeocete evolution (*i.e*., from 15–30 kg *Himalayacetus* [52.5 Ma] to 590–720 kg *Ambulocetus*/*Rodhocetus* [48 Ma]; 53-54). Body mass changes of similar scale followed the inactivation of *ucp1* in sirenians 46.0-44.8 Ma (Fig. 4A), the only other fully aquatic mammalian lineage. It is notable that loss of UCP1 function does not appear to be linked to the advent of aquatic birth in these lineages (pinnipeds and other semi-aquatic mammals are born on land or ice and generally do not enter the aquatic environment until after weaning) as our estimates of *ucp1* inactivation precede the loss of hindlimb land locomotion—and hence emergence of fully aquatic lifestyles—in primitive dugongids (55) and early basilosaurids (*e.g*., *Basilosaurus*, *Durodon*; 56) during the Bartonian by 5 to 10 Ma. Notably, absence of UCP1-mediated NST is not mechanistically linked with inferior thermoregulatory capabilities in neonates of either lineage as evidenced by year-round resident high-Arctic whales (*e.g*., bowhead, beluga, and narwhal) and the recently extinct sub-Arctic Steller’s sea cow (*Hydrodamalis gigas*). The presence of a pseudogenic copy of *ucp1* in woolly mammoths further precludes a UCP1-mediated thermogenic function for the BAT-like tissue described from a 1-month old specimen (24), and again illustrates that *ucp1* is not uniquely associated with the evolution of extreme cold hardiness in placental mammals.

In addition to abrupt changes in body mass (Fig. 4A), the inactivation of *ucp1* also coincided with rapid species diversification evident in the fossil record of these lineages (Fig. 4B), consistent with recent work linking increased body mass with cladogenesis (57). Notably, however, as most lineages lacking a functional UCP1 currently exhibit relatively low species diversities, we performed analyses with binary state speciation and extinction (BiSSE; 58) on an eutherian mammal species tree with 5138 taxa (see Materials and Methods; 59), as implemented in the diversitree package for R (60), to determine if diversification rates among extant placental mammals have been higher in taxa that retain a functional copy of this gene (‘UCP1-plus’) versus taxa that have a pseudogenic copy of this gene (‘UCP1-minus’). These analyses provide support for the hypothesis that diversification rates have been higher in UCP1-plus taxa than UCP1-minus taxa (table S4). An important caveat is that BiSSe is prone to high Type 1 error rates even for neutral characters that are not associated with trait-dependent diversification rates (61). We therefore simulated neutral characters on the same mammalian species trees that we employed for the above UCP1 diversification analyses, and confirmed that BiSSe has a high Type I error rate (p<0.05 in 43 out of 100 simulations) for our trees (table S5). However, 35 of these runs resulted in p values that were less supported than the least significant p value (0.000023) for UCP1.

The observation that brown-throated sloths (*Bradypus variegatus*) exposed to cold exhibit markedly improved temperature regulation abilities when pregnant (34) suggests the presence of inducible NST mechanism(s) in this and potentially other clades lacking a functional UCP1. One candidate recently implicated in muscle-based NST is sarcolipin (*sln*; 62, 63), which intriguingly exhibits two unique residue deletions in equids that may alter ATP-dependent Ca^2+^ uptake in skeletal muscle, though this gene inactivated in extant sloths (fig. S3). It is unlikely that other paralogs (*i.e*., *ucp2* and *ucp3*) assumed a UCP1-like role in BAT in these species as d*N*/d*S* analyses failed to identify differential selection acting upon these loci in branches where *ucp1* was lost (data files S4-S5).

Importantly, these ancillary analyses pinpoint additional pseudogenization events (*e.g*., *ucp2* and *ucp3* inactivation in armadillos [table S6]) that provide natural model systems to help elucidate the precise function of these proteins.

## DISCUSSION

Although NST is often thought to be an effective means to counter low ambient temperatures, heavy reliance on thermogenesis comes at a high energetic cost that may be difficult to meet in species that consume energy poor diets and exhibit marked reductions in metabolic intensity (xenarthrans and pangolins). Selection for larger body size is also predicted to reduce reliance on BAT by decreasing the relative surface area for heat loss (20, 21) and enabling the development of size-dependent heat conservation measures (*e.g*., countercurrent rete). In this regard, selection pressures contributing to the loss of *ucp1* may also be expected to favor precocial species due to their heightened insulation at birth and generally transient reliance on BAT-mediated NST relative to altricial taxa (64). In this context, it is notable that only one of the eight lineages lacking a functional *ucp1* locus is altricial (pangolins).

In light of the apparent link between our *ucp1* inactivation estimates and body mass changes evident in the fossil record (Fig. 4A), it is perhaps surprising that *ucp1* is intact and ostensibly evolving under purifying selection in large-bodied rhinoceros, pinnipeds (walrus, Weddell seals, harbor seals), and hippopotamuses (ω=0.25 to 0.29; table S2). This suggests that other factors including diet, metabolic intensity, developmental strategy, litter size, and birth weight, together with potential adaptive roles in brite/beige tissue, may have also played a role in the retention (or loss) of UCP1. Nonetheless, relatively low ω values for the above branches should be treated with caution in the absence of protein expression and functional data, as long branches may conflate multiple evolutionary histories (*i.e*., a recent transition to neutral selection that is obscured by a much longer period of purifying selection). Indeed, *ucp1* expression was not detected in ∼32 kg Weddell seal nor in ∼27 kg hooded seal neonates, both of which lacked conspicuous BAT depots (65); hippopotamuses similarly appear to lack BAT as newborns (17). Conversely, the intact *ucp1* locus of camels intriguingly exhibits a signature of neutral evolution (ω=1.03; table S2) coupled with a four-residue deletion within a highly conserved region of the third matrix loop that borders the GDP-binding domain, which may compromise its function (fig. S2).

The regressive evolution of *ucp1* together with attendant increases in body mass may thus have contributed to the initial exploitation of seasonal temperate niches (and other large-bodied terrestrial and aquatic habitats vacated by dinosaurs/marine reptiles following the KPg extinction) by these lineages and presumably underlies their rapid diversification apparent in the fossil record (Fig. 4B). This evolutionary success was, however, not devoid of negative long-term fitness consequences. For example, the re-evolution of small body size (<5 kg) is presumably constrained by competitive disadvantages versus species with UCP1-mediated NST, or has restricted medium-sized members of these lineages to tropical/subtropical environments (*i.e.*, xenarthrans, pangolins, hyraxes). These factors coupled with observations that large-bodied taxa exhibit higher risks of extinction (66) lead us to hypothesize that loss of UCP1-dependent thermogenesis may represent a historical contingency that underlies the current low species diversity of placental clades lacking a functional *ucp1* gene (Fig. 4B). This assertion is supported by our BiSSE analyses, as we demonstrate higher diversification rates in UCP1-plus taxa than UCP1-minus taxa. Although we confirm that this method is susceptible to high Type 1 error rates, most p values arising from our simulations were substantially less supported than those of our diversification analysis (*cf*. table S5 and table S6). Thus, to our knowledge, these results provide the first evidence that links diversification rates and hence species selection to the presence or absence of a single gene product. A second limitation to this interpretation is that other traits, which are unrelated to UCP1 inactivation, may also have contributed to lower diversification rates in UCP1-minus taxa. One possible example is the complete loss of teeth (edentulism), which occurs in four UCP1-minus clades (pangolins, anteaters, baleen whales, Steller’s sea cow) with highly specialized diets and feeding adaptations. All of these groups also have inactivating mutations in one or more tooth-specific genes (28, 67) that effectively preclude the reappearance of teeth in these taxa.

As adult and obese humans possess only negligible brown adipose depots, species lacking UCP1 provide unexplored model systems for adipocyte remodeling and alternative pathways of thermogenesis in response to cold that may be more transformative to biomedical applications. Taken together, our findings raise important questions regarding the roles and physiological significance of uncoupling proteins over the course of eutherian evolution, challenge the perception that BAT underlies cold tolerance in large-bodied eutherian mammals, and identify *ucp1* inactivation as a historical contingency that may underlie the current low species diversity of these clades.

### Materials and Methods

#### Taxon Sampling

This study included representatives of 141 vertebrates, 133 of which are mammals (1 monotreme, 4 marsupials, 4 xenarthrans, 11 afrotherians, 59 laurasiatherians, and 54 euarchontoglires). Outgroup representatives included 7 fish and 1 amphibian species. Due to the documented loss of uncoupling protein 1 (*ucp1*) in the archosaur lineage (*68*), birds and reptiles were not included in the dataset. We employed genome mining, polymerase chain reaction (PCR), and hybridization capture to obtain *ucp1*, *ucp2*, *ucp3*, and sarcolipin (*sln*) coding sequences examined in this study. Taxon sampling and GenBank accession numbers are provided in table S1.

#### Genome mining

We used key word searches to identify *ucp1, ucp2*, *ucp3*, and *sln* mRNA sequences in GenBank. We also performed standard nucleotide blasts (blastn; *69*) against whole-genome shotgun contigs using the “discontinuous megablast” setting and human *ucp1, ucp2*, *ucp3*, or *sln* coding sequences (accession numbers: NM_021833.4, U82819.1, U84763.1, NM_003063.2) as queries. Retrieved contigs from the top blast hits for each species were imported into the program Sequencher version 5.0, and manually annotated by aligning human *ucp/sln* exons against the individual contigs. In cases where putative exon lengths did not match those of the human loci, exon/intron boundary splice sites were identified following the GT-AG rule (*70*). For fragmentary and highly degraded *ucp1* pseudogenes (*e.g*., armadillo, pangolin, hyrax, sloth), we generated dot-plots versus *ucp1* mRNA sequences of closely related species using the EMBOSS:dotmatcher feature in Geneious R6.1 to identify exons. Linear comparison figures illustrating coding regions of these genes were prepared using Easyfig 2.1 (*71*).

The synteny of *ucp1* is highly conserved among vertebrates with *ucp1* located between the upstream *tbc1d9* and downstream *elmod2* loci (13, *68*). Thus, wherever possible, exons of these flanking genes were annotated on *ucp1*-containing contigs to ensure our annotated genes are *ucp1* orthologs. For example, a gene fragment (exon 6) housed on GenBank contig ABDV02364175.1 of Hoffman’s two-toed sloth (*Choleopus hoffmanni*) was on the same contig (7.6 kb upstream) from *elmod2*, and thus positively identified as *ucp1*.

Finally, discontiguous megablasts against the NCBI sequence read archive (SRA) database were performed for the woolly mammoth (*Mammuthus primigenius*), donkey (*Equus asinus*), fin whale (*Balaenoptera physalus*), and snow leopard (*Panthera uncia*) using queries of known *ucp* coding sequences from the nearest phylogenetic relative (listed below). Retrieved SRAs were imported into Sequencher and assembled against respective reference *ucp* contigs of the African elephant (*Loxodonta africana*), horse (*Equus caballus*), minke whale (*Balaenoptera acutorostrata*), and tiger (*Panthera tigris altaica*) to generate consensus *ucp1*, *ucp2*, *ucp3*, and *sln* coding sequences.

#### DNA amplification and Sanger sequencing

The African elephant *ucp1* sequence on GenBank exhibited a 1 bp frameshift deletion in exon 3 (incorrectly annotated on Ensembl). We thus verified this deletion in both African and Asian elephant (*Elephas maximus*) DNA samples via PCR and Sanger sequencing techniques. PCR was also used to obtain *ucp1* sequences from the three-toed brown-throated sloth (*Bradypus variegatus*), silky anteater (*Cyclopes didactylus*), bowhead whale (*Balaena mysticetus*), Grant’s zebra (*Equus quagga boehmi*), black rhinoceros (*Diceros bicornis*), and Indian rhinoceros (*Rhinoceros unicornis*). Specimen information is provided in table S7 (67, *72*-*74*).

Genomic DNA extractions were performed using the DNeasy Blood and Tissue Kit (Qiagen). For the bowhead whale sample, a whole genome amplification was performed using a REPLI-g Mini Kit (Qiagen) following the manufacturer’s directions prior to PCR amplification to increase the amount of available DNA. Primers were designed to target each of the six individual *ucp1* exons from the available draft genome of the nearest phylogenetic relative using the Primer Premier 5.0 software. PCRs were performed using 2 units of One*Taq* DNA polymerase (New England Biolabs) in 20 μL reactions using the following thermal cycling profile: initial denaturation at 94°C for 2 min 30 sec, followed by 35 cycles at 94°C (denaturation) for 30 sec, 48-62°C (annealing) for 30 sec and 68°C (extension) for 30 sec, followed by a final extension of 5 min at 68°C. Products were visualized on 1.5% agarose gels. In cases where this protocol did not result in a distinct amplification product, nested or hemi-nested PCRs were performed using aliquots of the initial amplification reactions as template DNA. Distinct bands of appropriate size were excised from agarose gels and purified using a GeneJet Gel Extraction Kit (Fermentas). The purified PCR products were either sequenced directly or cloned into pDdrive cloning vectors (Qiagen) and transformed into *Escherichia coli* plasmids (New England Biolabs). Clones were grown overnight on agar plates at 37°C. Positive clones were identified using a blue-white screening technique, followed by PCR amplification and visualization of the targeted DNA by gel electrophoresis. Cells containing inserts of the correct size were grown overnight in Luria broth culture medium and the DNA was subsequently purified using the Zyppy Plasmid Miniprep Kit (Zymo Research). Sequencing reactions were conducted using the BigDye Terminator v3.1 Cycle Sequencing Kit (Applied Biosystems) as per the manufacturer’s directions. Sequencing reactions were purified using the Zymo Research DNA Sequencing Clean-up Kit (Zymo Research) and sequenced in both directions with an Applied Biosystems 3130 Genetic Analyzer.

#### DNA hybridization capture and next-generation sequencing

Methods for obtaining *ucp1*, *ucp2*, *ucp3*, and *sln* sequences from the dugong (*Dugong dugon*) and recently extinct Steller’s sea cow (*Hydrodamalis gigas*) are detailed elsewhere (67, *73*). Briefly, barcoded DNA libraries suitable for Illumina sequencing were prepared from dugong and Steller’s sea cow DNA extracts and hybridized to Agilent SureSelect Capture arrays imprinted with the coding region (plus 25-30 bp of flanking sequence for each exon) of African elephant *ucp1*, *ucp2*, *ucp3*, and *sln* sequences. Eluted DNA fragments were sequenced on Illumina GAIIx and HiSeq2500 genome analyzers, trimmed of adapter sequences, and those <20 bp removed from the dataset (67). Remaining reads were assembled to manatee reference sequences using Geneious. The Steller’s sea cow assemblies were manually examined for C→U[T] and G→A DNA damage artifacts (*75*), with any polymorphic C/T or G/A sites subsequently scored as C or G; non-polymorphic C→T or G→A changes relative to dugong or manatee sequences were treated as genuine (67).

For pygmy hippopotamus (*Choeropsis liberiensis*) and beluga (*Delphinapterus leucas*), genomic DNA was extracted using the DNeasy Blood and Tissue kit (Qiagen). Three μg of genomic DNA was sheared into fragments of highest concentration at ∼180 to 190 bp using a Bioruptor (Diagenode), followed by treatment with PreCR Repair Mix (New England Biolabs). The SureSelect ^XT^ Target Enrichment System for Illumina Paired-End Sequencing Library kit (Agilent) was used for library construction and target enrichment. Target enrichment was done with a custom-designed SureSelect biotinylated RNA library that included cow *ucp1*, *ucp2*, and *ucp3* sequences. Target enriched libraries were paired-end 2x100 sequenced on an Illumina HiSeq 2500 platform at the UC Riverside Institute for Integrative Genome Biology Genomics Core. Per base quality distributions of de-multiplexed fastq files were visualized for both read pair files using FastQC v.0.10.0 (http://www.bioinformatics.babraham.ac.uk/projects/fastqc/) with the no group setting. Based on these results, FASTX-Toolkit v.0.0.13.2 (http://hannonlab.cshl.edu/fastx_toolkit/index.html) was used to trim the first 3 bases and the last base, which resulted in 97 bp reads. Reads with all but three identical bases or a quality score below 30 at any base position were then filtered out. PRINSEQ lite v.0.20.4 () was then used to find read pairs that passed the filtering conditions. These read pairs were then interleaved into a single file using the ShuffleFastq script in RACKJ v.0.95 (http://rackj.sourceforge.net/Scripts/index.html#ShuffleFastq).

DNA libraries were also constructed for the Malayan tapir (*Tapirus indicus*), black rhinoceros (*Diceros bicornis),* Indian rhinoceros (*Rhinoceros unicornis*), Sumatran rhinoceros (*Dicerorhinus sumatrensis*) and the extinct woolly rhinoceros (*Coelodonta antiquitatis*). Briefly, previously extracted genomic DNA samples of the black and Indian rhinoceroses (see above) were amplified using a REPLI-g Mini Kit (Qiagen) to increase the quantity of available DNA. DNA fragments were then enzymatically sheared using NEBNext dsDNA Fragmentase (New England Biolabs) and libraries were created using NEXTFlex barcoded adaptors and a NEBNext Fast DNA Library Prep Set for Ion Torrent (New England Biolabs) following the manufacturer’s protocol. A DNeasy Blood and Tissue Kit (Qiagen) was used to perform extractions from Malayan tapir and Sumatran rhinoceros blood samples, while ancient DNA was extracted from three woolly rhinoceros bone samples in an ancient DNA dedicated laboratory following the methods described by Dabney *et al. (77*). DNA libraries were constructed for these species using a NEBNext DNA Library Prep Master Mix Set for 454 (New England Biolabs). All libraries were then size selected using an E-gel iBase (Invitrogen) and target enriched using biotinylated 120mer MyBaits RNA probes (Mycroarray) designed with a 4x tilling pattern from the draft genome of the white rhinoceros (*Ceratotherium simum*) following the manufacturer’s protocol. Enriched DNA libraries were then amplified in 25 μl PCR cocktails with the following thermocycling profile: initial denaturation 98°C for 30 sec, 14-20 cycles of 98°C for 20 sec (denaturation), 58°C for 30 seconds (annealing), and 72°C for 30 sec (extension) followed by a final extension period of 72°C for 5 min. Libraries were sequenced at the University of Manitoba on an Ion Torrent PGM platform using an Ion PGM Hi-Q Sequencing Kit and Ion 314 v2 and Ion 318 v2 barcoded chips (Applied Biosystems). Sequenced reads were assembled to white rhinoceros *ucp1*, *ucp2*, *ucp3*, and *sln* reference sequences in Geneious R7.1.9 at 20% maximum mismatch to build consensus sequences. Ancient DNA reads from the woolly rhinoceros libraries were examined for C→T and G→A mutations as described above for the Steller’s sea cow.

#### UCP coding sequence alignments

Translated *ucp* coding sequence files were created for each species and all sequences visually examined for nonsense, insertion or deletion frameshift, start codon, termination codon, and splice site mutations. Multiple nucleotide sequence alignments were constructed for *ucp1*, *ucp2*, and *ucp3* datasets, both individually and combined, using the program MUSCLE 3.6 (*78*), and manually adjusted to eliminate ambiguous regions. The single *ucp1* sequence retrieved from the SRA database of the Darwin’s ground sloth (*Mylodon darwinii*) was deemed to be too short for selection pressure analyses and therefore excluded from the final *ucp1* alignment. Insertions unique to fish (9 bp), marsupials (12 bp), and *Canis lupus familiaris* (6 bp) were also excluded from the *ucp1* alignment. The final *ucp1* alignment included 141 species and totaled 936 bp in length (data file S6). The *ucp2* alignment included 131 species and totaled 948 bp in length (data file S7), the *ucp3* alignment included 128 species and totaled 936 bp in length (data file S8), while the combined *ucp1*-*ucp2*-*ucp3* alignment included 400 loci and totaled 983 bp in length.

#### UCP gene tree analysis

To further ensure all *ucp* sequences were correctly assigned, a maximum likelihood *ucp1*-*ucp2*-*ucp3* tree was constructed with the RAxML 7.2.8 plugin in Geneious using the GTR + Γ option (fig. S1). The *ucp* tree was generated starting with a randomized tree and 500 bootstrap replicates using the “rapid bootstrapping setting and search for the best-scoring ML tree” parameter.

#### Species tree analysis

We constructed a species tree for 136 mammals and four outgroups (western clawed frog [*Xenopus tropicalis*], Carolina anole [*Anolis carolinensis*], red junglefowl [*Gallus gallus*], and zebrafinch [*Taeniopygia guttata*]) on the basis of a supermatrix for 30 nuclear (*A2AB*, *ADRB2*, *APP*, *ATP7A*, *Adora3*, *ApoB*, *BCHE*, *BDNF*, *BMI1*, *BRCA1*, *BRCA2*, *CHRNA1*, *CMYC*, *CNR1*, *CREM*, *DMP1*, *ENAM*, *EDG1*, *FBN1*, *GHR*, *IRBP*, *MC1R*, *PLCB4*, *PNOC*, *Rag1*, *Rag2, SWS1*, *TTN*, *TYR1*, *VWF*) and 11 mitochondrial genes (*12S rRNA*, *16S rRNA*, *CYTB*, *COI*, *COII*, *COIII*, *ND1*, *ND2*, *ND3*, *ND4*, *ND5*). Sequences were obtained from GenBank, Ensembl, PreEnsembl, and other genomic resources (*i.e*., gigadb.org for *Ursus maritimus* and *Pantholops hodgsonii*, www.bowhead-whale.org for *Balaena mysticetus*) and were culled from larger sequence alignments for all available mammalian species (M.S.S., unpublished). Accession numbers, scaffold numbers, etc., are given in data file S9. Alignment-ambiguous regions were excluded prior to phylogenetic analysis. Additional sites with missing data/gaps for our reduced set of species were removed in RAxML. The final alignment comprises 50,879 bp (data file S10). A species tree was obtained with RAxML-HPC2 on XSEDE (RAxML version 8.1.11) on CIPRES (*79*). The RAxML analysis was performed with 32 partitions, each of which was given its own GTR + Γ model of sequence evolution. The 32 partitions included one for each of 30 nuclear genes, one for 12S rRNA + 16S rRNA, and one for nine mitochondrial protein-coding genes. We employed the GTRGAMMA model for both rapid bootstrap analysis (500 bootstrap iterations) and a search for the best ML tree.

#### Timetree analyses

Timetree analyses were performed on the 41-gene supermatrix tree with the mcmctree program in PAML 4.5 (27), which implements the MCMC algorithms of Rannala and Yang (*80*). Analyses were performed with the autocorrelated rates model. Each of 32 partitions (see above) was allowed to have its own GTR + Γ model of sequence evolution. We set one time unit = 100 million years (Ma). Analyses were run with cleandata = 0. Shape (α) and scale (β) parameters for the gamma prior of the overall rate parameter μ (*i.e*., rgene_gamma in mcmctree) were 1 and 4.74, respectively. Calculations for the shape and scale parameters of the gamma prior for the rate-drift parameter (*i.e*., sigma2_gamma in mcmctree) assumed an age of 344 Ma for the most recent common ancestor of Tetrapoda (average of minimum and maximum constraints in Benton *et al*.; *81*). Chains were run for 100,000 generations after a burn-in of 10,000 generations, and were sampled every 20 generations. We employed hard-bounded constraints for 37 nodes including the root node (Tetrapoda). Minimum ages were based on the oldest crown fossils that are assignable to each clade while maximum ages were based on stratigraphic bounding, phylogenetic bracketing, and phylogenetic uncertainty (49, 67, *82*, *83*). Stratigraphic bounds were extended by one stage for younger deposits (late Miocene, Pliocene, Pleistocene) and two stages for older deposits (middle Miocene and earlier). Minimum and maximum constraints (56, 67, *81*, *84*-*165*) are summarized in table S8. Stage boundaries are from the International Chronostratigraphic Chart v 2014/02 (www.stratigraphy.org; *166*).

#### Evolutionary selection pressure and inactivation analyses

The CODEML program in the PAML 4.8 software package (27) was employed to estimate the nonsynonymous/synonymous substitution rate ratio (d*N*/d*S* or ω value) in order to infer the modes of natural selection acting upon the three *ucp* loci. All frameshift insertions in the alignments were deleted, and nonsense mutations were recoded as missing data prior to analysis. Our constructed species tree was used to guide the phylogeny of all CODEML runs with outgroup fish species added to the guide tree following the phylogenetic relationships described by Near *et al*. (*167*). All CODEML runs were performed using the F3x4 codon frequency model. The free-ratio (M1) model, which calculates an independent ω value for every branch of the tree, was performed for all three *ucp* alignments (data files S6-S8). Results of this analysis for *ucp1* were used as a starting point to target specific lineages later tested with the branch (M2) model for neutral evolution or positive selection. Target lineages for these analyses included the stem placental branch, the stem pinniped branch, the stem rhinoceros branch, the pygmy hippo branch, the camel branch, and finally all pseudogenic *ucp1* branches. Individual M2 branch models for the stem placental, pinniped, rhinoceros, pygmy hippo, and camel branches were performed using the same branch categories described below for the *ucp1* pseudogene analysis plus an additional branch category to target the lineage(s) of interest. Likelihood ratio tests (table S2) were then performed against the *ucp1* pseudogene M2 branch model to test if the obtained ω values were significantly different from all functional placental branches. The M2 models were run with the method = 0 setting which calculates all branch lengths simultaneously.

Branch-site analyses were also performed to identify potential amino acids under positive selection that may have contributed to the gain of non-shivering thermogenic function of *ucp1* in a stem placental ancestor. Here, the stem placental branch of the species tree was set to the foreground while all others were background branches. To reduce the possibility of signatures of neutral evolution skewing the results, pseudogenes were not included in this analysis, resulting in a data set of 115 species. The branch-site MA model, under the Bayes empirical Bayes method, resulted in the identification of eight sites with a posterior probability >0.95 of being greater than one (table S2). However, when tested against the null model, where the ω values of positively selected sites under the MA model are fixed at neutrality (ω = 1), the likelihood ratio test demonstrated that omega values for these sites were not significantly greater than one (2Δl = 0.706, df = 1, p = 0.400).

For each *ucp1* pseudogenization event we calculated the transition point from purifying selection to neutral evolution following the method and equations of Meredith *et al*. (28). The *ucp1* inactivations are considered to have occurred at or following this date. Branches were first divided into three categories: functional, transitional (mixed), and pseudogenic. Functional branches are those that lead to internal or terminal nodes with an intact copy of *ucp1* and presumably evolved under functional constraints. Transitional branches record the first evidence of *ucp1* pseudogenization (*e.g*., stop codon, frameshift mutation) and are deemed to contain both functional and pseudogenic segments that evolved under purifying selection and neutral evolution, respectively. Pseudogenic branches post-date mixed branches and are expected to have *dN*/*dS* values at or near the neutral value of 1 in the absence of purifying or positive selection (28). In taxonomic clades where all members share inactivating *ucp1* mutations (*i.e*., Cetacea, Equidae, Proboscidea, Sirenia, Pilosa), the transitional branches were deemed to be immediately ancestral to the radiation of the clade. The guide tree was labeled in order to calculate ω values for all functional non-mammalian vertebrate branches, all functional non-placental mammalian branches, all functional placental mammal branches, and all pseudogenic placental mammal branches as distinct branch categories as well as each individual transitional branch using the M2 branch model. The equations developed by Meredith *et al*. (28) were employed to estimate transition dates from purifying selection to neutral evolution of *ucp1* along transitional branches using our upper and lower nodes for each branch from our time-calibrated phylogeny (data file S1).

#### Paleotemperature calculations

We used stable oxygen isotope (δ^18^O) values (‰) of benthic foraminifera (supplementary Fig. 3 dataset of Friedrich *et al*.; *29*) as a proxy for global temperature over the last ∼82 Ma (see Fig. 3). Within this window, paleotemperatures were calculated (30) from the Cretaceous to Oligocene as: temperature (°C) = 16.5-4.8(δc-δw), where δc is the measured δ^18^O value of foraminifera calcite and δw (-1.0 ‰) is the estimated value for an ice-free ocean (31).

#### Fossil body mass and diversity

Fossil body mass estimates recorded at each geological stage for Cetacea, Sirenia, Proboscidea, Hyracoidea, and Equidae were assembled from the literature (44, 53, 54, *168*-*184*) (data file S3). We also downloaded all species occurrence records for these eutherian clades from each geological stage ranging from the Selandian to the Holocene from the Paleobiology Database (www.fossilworks.org) and other literature sources (44, 46, *168*, *175*, *182*, *185*-*197*). Species and genera counts were then plotted for the median date of each stage (data file S3). Owing to fragmentary fossil records and few body mass estimates for early Xenarthra, Pholidota, and Suidae, these groups were not included in these analyses.

#### Diversification analysis

We selected five fully resolved timetrees (numbers 1, 101, 201, 301, 401) from Faurby and Svenning (59) to allow for phylogenetic uncertainty. Each of these trees included all extant mammalian species plus late Quaternary extinct mammals. We pruned all non-placental taxa from these trees. For the remaining placentals, we only retained extant taxa and recently extinct taxa that are included in Wilson and Reeder (*198*). After pruning, our trees comprised 5138 placental species. The presence versus absence of a functional copy of *ucp1* was inferred from the condition of the gene in the closest relative for which there is DNA sequence evidence, supplemented by evidence from shared mutations and our estimates for inactivation times of *ucp1* in UCP1-minus lineages. For example, molecular data provide evidence for inactivation of *ucp1* in representatives of all three clades of Xenarthra (armadillos, anteaters, sloths), and our inactivation date for this gene is estimated in the latest Cretaceous. We therefore coded all xenarthrans as ‘UCP1-minus’. Similarly, diverse rodent and bat taxa with genome sequences all have intact copies of *ucp1*. Based on this evidence, and the general absence of very large rodents and bats, we coded all extant rodent and bat species as ‘UCP1-plus’. Data file S3 contains all of the inferred character states for *ucp1* that were employed in our diversification analyses, which were performed with the BiSSE method (58) as implemented in diversitree in R (60). We compared two diversification models, each of which fixed the transition rate from UCP1-minus to UCP1*-*plus at zero given the absence of empirical evidence for such transitions and the low probability of resuscitating a functional copy of a gene from an inactivated copy with frameshift and/or premature stop codon mutations. The first model allowed for separate speciation (lambda) and extinction (mu) rates in UCP1-plus taxa versus UCP1-minus taxa, whereas the second model constrained equal diversification rates in UCP1-plus taxa and *UCP1*-minus taxa (*i.e*., lambda0 = lambda1, mu0 = mu1). Simulations with neutral characters were performed with the same five trees from Faurby and Svenning (59). We performed 20 simulations for each of these trees (total = 100) and used the same q01 transition rates that were inferred from analyses with the UCP1 data (table S4).

## SUPPLEMENTARY MATERIALS

Supplementary material for this article is available at http://advances.sciencemag.org/ Materials and Methods

fig. S1. Schematic maximum-likelihood tree of *ucp* sequences used in this study (N=400).

fig. S2. Exon alignments depicting deleterious mutations found in *UCP1* sequences of placental mammal taxa.

fig. S3. Ribbon diagram of residues 13 to 304 of human UCP1 (UniProt accession number P25874) structurally modelled by SWISSMODEL.

fig. S4. Amino acid alignment of vertebrate sarcolipin.

table S1. GenBank accession numbers of species used in this study.

table S2. Likelihood ratio tests for *ucp1* CODEML models.

table S3. Calculated *ucp1* inactivation dates (millions of years ago; Ma) within the placental mammal lineage.

table S4. Results of binary state speciation and extinction (BiSSE) models with and without constraints on the diversification rate for five trees from Faurby and Svenning (*194*).

Table S5.

table S6. Deleterious mutations found in *ucp2* and *ucp3* sequences of placental mammal taxa.

table S7. Specimen data and sources of tissue samples used for PCR amplification and DNA hybridization capture.

table S8. Fossil constraints used in timetree analysis.

Data file S1. Gene partitions, a maximum likelihood phylogram, and mcmctree [text file].

Data file S2. Uncoupling protein 1 (*ucp1*) free ratio model [text file].

Data file S3. Fossil body mass, taxon diversity, and UCP1 diversification codings [Excel file].

Data file S4. Uncoupling protein 2 (*ucp2*) free ratio model [Text file].

Data file S5. Uncoupling protein 3 (*ucp3*) free ratio model [Text file].

Data file S6. Uncoupling protein 1 (*ucp1*) coding sequence alignment [Fasta file].

Data file S7. Uncoupling protein 2 (*ucp2*) coding sequence alignment [Fasta file].

Data file S8. Uncoupling protein 3 (*ucp3*) coding sequence alignment [Fasta file].

Data file S9. Accession numbers for the 41 loci used to construct the 51 kb maximum likelihood species tree [Fasta file].

Data file S10. 51 kb alignment (41 genes) for 140 vertebrate species [Text file].

## Acknowledgments

We thank Stephen Petersen, John Gatesy, Cheryl Y. Hayashi, Ross D.E. MacPhee, Eske Willerslev, Mads Bertelsen, R. Havmøller, and Tom Gilbert for providing tissue samples, Oliver Friedrich for assistance with δ^18^O isotope to temperature conversions, and Mark Uhen and Felisa Smith for providing access to the MAMMOTH v1.5 database. Authorization to use paintings by Carl Buell was provided by John Gatesy.

## Funding

This work was supported by NSERC Discovery Grants to K.L.C. (238838) and J.R.T. (418503), an NSERC Discovery Accelerator Supplement to K.L.C. (412336), the Canada Research Chairs program to J.R.T. (950-223744), a German Center for Diabetes Research (DZD) grant to M.J., and a NSF grant (DEB-1457735) to M.S.S. M.J.G. was funded in part by a University of Manitoba Undergraduate Research Award, a University of Manitoba Graduate Fellowship, and a Manitoba Graduate Fellowship. J.L.A.P. was funded in part by a Natural Environment Research Council (NERC) student fellowship and the University of York. A.V.S. was funded in part by a National Sciences and Engineering Research Council (NSERC) Alexander Graham Bell Canada Graduate Scholarship–Doctoral Program. **Author contributions:** M.J.G. conceived the project, designed research, conducted experiments, performed selection and dating analysis, interpreted the results, prepared the figures, and assisted with manuscript writing; K.L.C conceived the project, designed research, interpreted the results, prepared the figures, and drafted the manuscript; M.J., J.R.T., and M.H. designed research and interpreted the results; J.S., J.L.A.P., N.W., and A.V.S. conducted the hybridization capture and sequencing experiments; M.S.S. performed phylogenetic, timetree, selection, and diversification analysis, interpreted the results, and assisted with manuscript writing; All authors discussed the results and helped revise the manuscript. **Competing interests:** The authors declare that they have no competing interests. **Data Availability:** New sequences obtained in this study are archived at GenBank (accession numbers XXX to YYY). All data needed to evaluate the conclusions in the paper are present in the paper and/or the Supplementary Materials. Additional data related to this paper may be requested from the authors.

